# Global Age-Specific Patterns of Cyclic Gene Expression Revealed by Tunicate Transcriptome Atlas

**DOI:** 10.1101/2020.12.08.417055

**Authors:** Yotam Voskoboynik, Aidan Glina, Mark Kowarsky, Chiara Anselmi, Norma F Neff, Katherine J Ishizuka, Karla J Palmeri, Benyamin Rosental, Tal Gordon, Stephen R Quake, Irving L Weissman, Rachel Ben-Shlomo, Debashis Sahoo, Ayelet Voskoboynik

**Affiliations:** Stanford University, Hopkins Marine Station, Pacific Grove, CA 93950, USA; Department of Physics, Stanford University, CA 94305, USA; Institute for Stem Cell Biology and Regenerative Medicine, Stanford University School of Medicine, Stanford, CA 94305, USA; Chan Zuckerberg Biohub, San Francisco CA 94158, USA; The Shraga Segal Department of Microbiology, Immunology, and Genetics, Faculty of Health Sciences, Center for Regenerative Medicine and Stem Cells, Ben-Gurion University of the Negev, Beer-Sheva, 84105, Israel; Zoology Department, Tel Aviv University, Tel-Aviv, 69978, Israel; Departments of Applied Physics and Bioengineering, Stanford University, CA 94305, USA; Ludwig Center, Stanford University School of Medicine, Stanford, CA 94305, USA; Department of Biology and Environment, University of Haifa-Oranim, Tivon 36006, Israel; Department of Pediatrics, University of California San Diego, La Jolla, CA, USA; Department of Computer Science and Engineering, Jacob’s School of Engineering, University of California San Diego, La Jolla, CA, USA

**Author notes:** These authors contributed equally to this work (Yotam Voskoboynik, Aidan Glina, Mark Kowarsky). Corresponding Authors E-mail: Ayelet Voskoboynik Debashis Sahoo Rachel Ben-Shlomo.

## Abstract

Expression levels of circadian clock genes, which regulate 24-hour rhythms of behavior and physiology, have been shown to change with age. However, a study holistically linking aging and circadian gene expression is missing. Using the colonial chordate *Botryllus schlosseri*, we combined transcriptome sequencing and stem cell-mediated aging phenomena to test how circadian gene expression changes with age. This revealed that *B. schlosseri* clock and clock-controlled genes oscillate organism-wide, with daily, age-specific amplitudes and frequencies. These age-related, circadian patterns persist at the tissue level, where dramatic variations in cyclic gene expression of tissue profiles link to morphological and cellular aging phenotypes. Similar cyclical expression differences were found in hundreds of pathways associated with known hallmarks of aging, as well as pathways that were not previously linked to aging. The atlas we developed points to alterations in circadian gene expression as a key regulator of aging.

**One Sentence Summary:** The Ticking Clock: Systemic changes in circadian gene expression correlates with wide-ranging phenotypes of aging

---

Circadian clocks are systems that oscillate with a consistent phase and control daily functions at the cellular, tissue, and organismal levels. Consisting of a small number of core clock genes operating within transcription-translation feedback loops, they activate and repress the expression of hundreds of downstream genes and regulate the response of cells to daily and seasonal fluctuations in light and temperature *(1–2)*. Proper functioning of the circadian rhythm system is essential for maintaining cellular and tissue homeostasis. The robustness of this system is reduced with age, resulting in disturbed circadian rhythms in old organisms relative to young adults *(3)*. This disrupted circadian control leads to disparities in the rhythms of waking activity, core body temperature, suprachiasmatic nucleus (SCN) firing, release of hormones such as melatonin and cortisol, and more *(4)*. Growing molecular evidence indicates that the circadian clock and aging process are closely associated *(3–8)*. The reduction in the expression levels of the core clock genes *Bmal1* and *Per2* in the mammalian SCN has been found to be age-associated *(5)*. The expression patterns of the other core clock genes in the aged mammalian SCN is less clear. Reports have indicated unchanged expression levels, reduced expression levels, or period lengthening *(7, 9)*. Changes in the expression patterns of the core clock in both aged and peripheral tissues have been inconsistent as well, with different species and even different tissues showing varying expression profiles *(4, 7, 8)*. Despite these inconsistencies, both aging and circadian rhythms are embedded in every tissue and organ of the body. Therefore, they must operate on the organismal level at some capacity. For this reason, we expect the molecular changes throughout the day due to age to be detectable on a systemic level. However, limitations of sampling and processing whole body tissues over time and age in conventional aging models make it difficult to directly monitor and study the important effect of changes in circadian rhythms on aging phenotypes.

Colonial chordates like *Botryllus schlosseri* provide a key to bridge this gap. This marine organism belongs to the chordate subphylum Urochordata, a taxonomic group considered to be the closest living invertebrate relative of vertebrates *(10)*. This group shares homology with at least 75% of the human gene repertoire *(11)*.

*B. schlosseri* exhibits several characteristics that render it a valuable model for the study of the impact of changes in circadian rhythms on aging. These include: (i) A long life span, with colonies in our lab living up to 20 years; (ii) Clonal asexual reproduction (Fig. S1), which allows separation of one genotype (colony) into several clonal replicates; (iii) Weekly stem cell-mediated asexual tissue regeneration of all body organs *(12–16;* Fig.S1), allowing for the study of stem cell aging’s effect at the organism level; (iv) Easy, serial collection of whole-body samples along their entire lifespan, which offers a unique approach to characterize genetic changes across time and age. Using *B. schlosseri*, this study sets out to holistically characterize the link between aging and circadian rhythms at the system, tissue, and organism level. Aiming to link organism-wide aging phenotypes and specific changes in circadian gene expression, we first **characterized***B. schlosseri* **aging and circadian behaviors**. Morphological phenotypic changes observed in young versus old *B. schlosseri* colonies include changes in pigmentation level (transparent vs. highly pigmented), terminating blood vessel (ampullae) shape (oval vs. elongated), significantly reduced zooid size as measured by average zooid area, and decreased regeneration capacity which wanes with age (measured by number of individuals (zooids)) in a colony (Fig. 1A-C; S1D). To further identify potential cellular changes that characterize the biology of aging in *B. schlosseri*, we compared the proportion of 24 cell populations (as described in *15*) between 3-5 days old (n=190) and 4-13 years old colonies (n=37) (Fig. 1E). This comparison revealed age-associated alterations in the cellular composition. Juveniles showed a higher level of enriched hematopoietic and germline stem cell populations compared to the higher level of engulfing phagocytic, cytotoxic, and pigment cells in aged colonies (Fig. 1E). We also found diurnal, circadian behaviors that show reduced nocturnal activity. Slower heart rate and siphon activity measured at night suggest that *B. schlosseri* is a diurnal organism. Young and mid-aged colonies demonstrated slower heartbeats at night (Fig. 1D) as well as infrequent siphon contractions (Movie S1) when compared to the day. Circadian changes in heart rate and siphon activity were not observed in aged colonies, demonstrating diurnal phenotypes diminishing with age. (Fig. 1D; Movies S1).

**Fig. 1.**
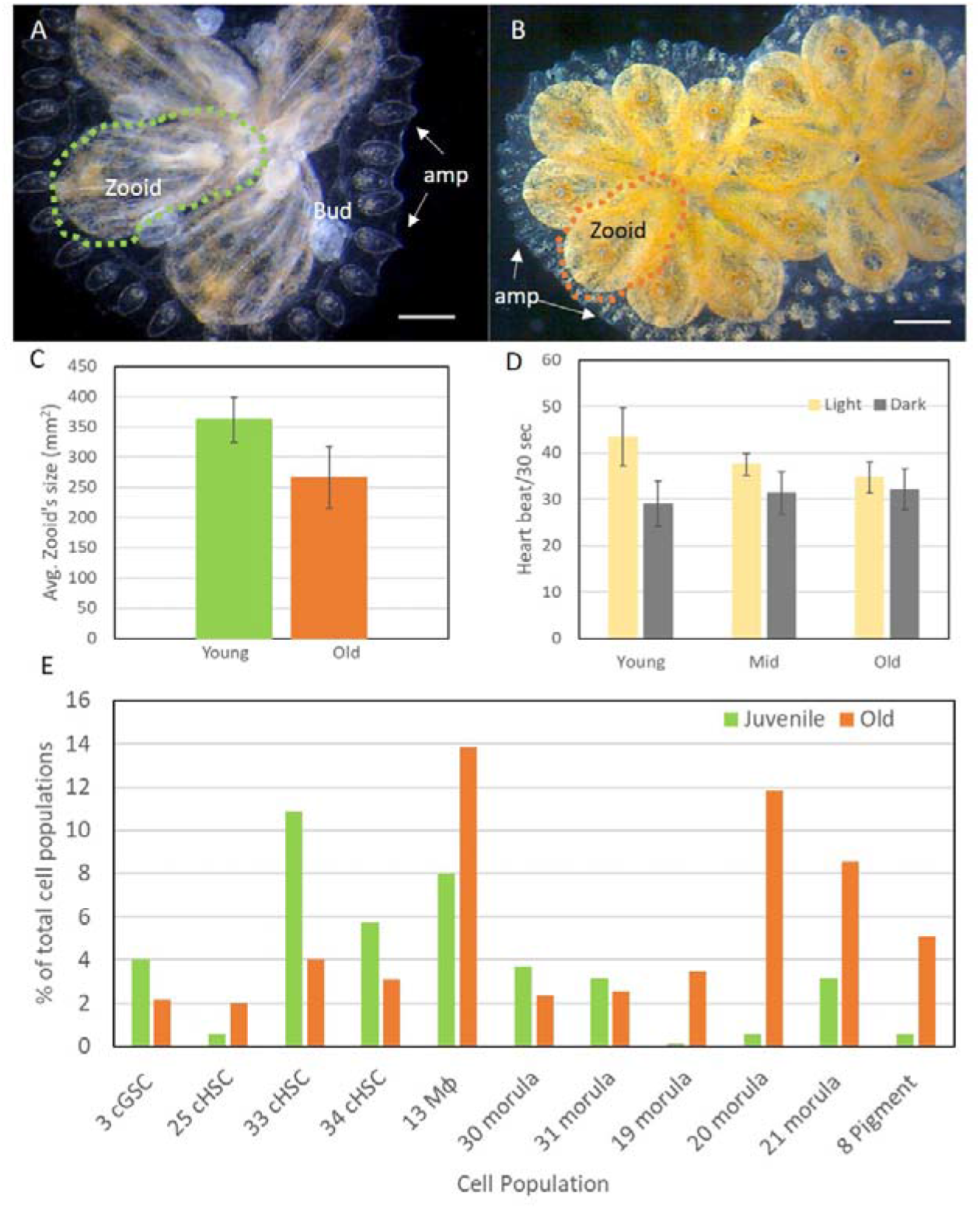
*B. schlosseri* aging phenotypes and diurnal behavior (A-B) Morphological changes observed in young (**A**) versus old (**B**). Zooid size (large vs. small), blood vessel/ampullae shape (oval vs. elongated), and pigmentation level (transparent vs. highly pigmented) are the main phenotypic changes observed with age. Scale bar-1mm, amp-ampullae, dotted circle outline zooid. (**C**) Zooid size is significantly reduced with age as measured by average zooid area (mm^2^). (**D**) We monitored colonies from 3 age groups during day (light) and night (dark) and discovered that the heartbeats of young (40 days, light n=12, dark n=10) and mid-age (1515 days old, light n= 9, dark n=6) are significantly slower at night. Young colonies averaged 43.412±6.25/30sec during the day compared to 29±4.8/30sec at night; mid-age colonies average heartbeat 37.44 ±2.35/30sec during the day, compared to 31.33±4.5/30sec at night. The difference between average heartbeat during day and night in the old group (6905 days old) is not significant, 34.55±3.35/30sec during the day, compared to 32.11±4.45/30sec at night. (**E**) Comparison between juvenile (3-5 days old, n=190) and old (4-13 years old, n=37) colonies revealed age-associated alterations in the frequency of specific cell populations. Juvenile colonies exhibited a higher frequency of enriched hematopoietic and germline stem cell populations (3, 33, 34), while older colonies showed a high frequency of engulfing, phagocytic, cytotoxic, and pigment cell populations (13, 19, 20, 21, 8). Cell population analysis was done using BD FACS Aria as described in *(15)*.

Next, we investigated the *B. schlosseri* genome (*11*), to **identify its circadian photoreceptors and core clock genes**(Table S1A; Fig. 2, S2). Three homologs of the vertebrate photoreceptor pinopsin/melanopsin were found. Homologs for core clock genes including multiple genes for βHLH - PAS domain transcription factors (*clock/bmal/arnt*) and multiple genes for clock regulators like casein kinase (ε and δ), Rev-erb, RorB, Glycogen synthase kinase 3 (*GSK3*; also known as Shaggy (*sgg*)), Timeless (*tim)*, Timeout, *Sirt*, and *Mtor*, were also identified. These represent a mix of genes resembling either vertebrate or invertebrate sequences (based on sequence homology; Table S1A). However, orthologs for the core clock suppressors *period* and *cryptochrome* were not found in the *B. schlosseri* genome assembly, as well as in 14 other tunicates genomes sequenced to date (aniseed database; *17*).

**Fig. 2.**
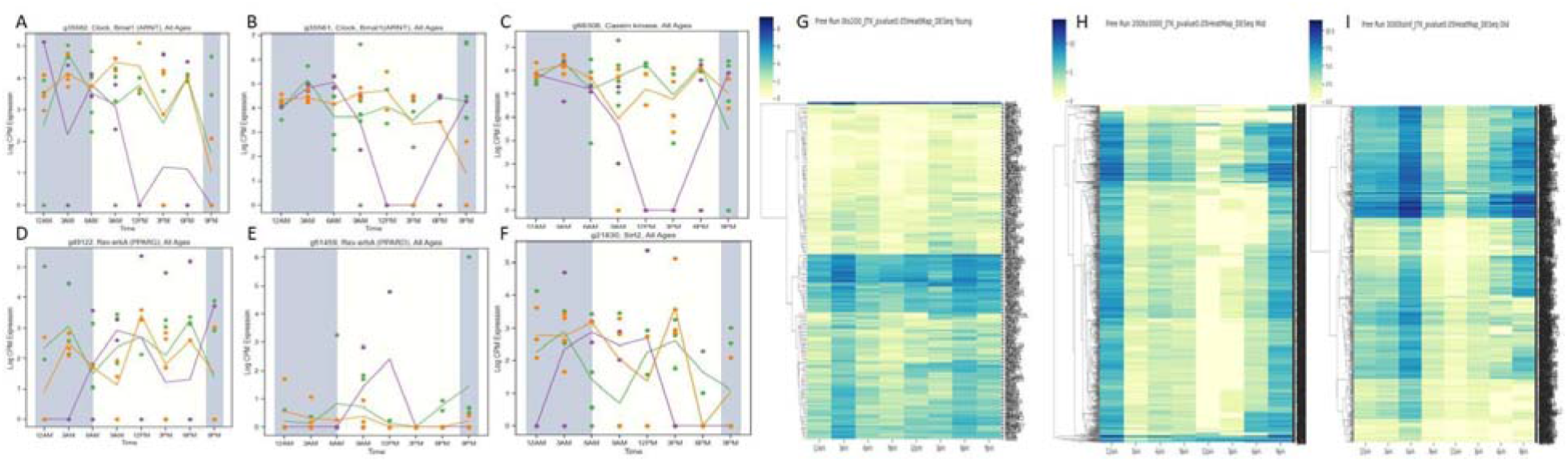
*B. schlosseri* circadian gene expression changed dramatically with age. **(A-F)** Age-associated changes in the cyclic expression of *B. schlosseri* putative clock and clock regulators. (**A-B**) In two putative *clock/bmal* genes that carry the bHLH-PAS domain, the mid-age group demonstrates large cycling patterns that are reduced in young and old colonies (**C**) Casein Kinase cycling patterns across all age groups resemble the cycling pattern of *clock/bmal* in plot B. (**D-E**) Two putative *Rev-erba* genes demonstrate opposite expression trends to *clock/bmal* cycling patterns, a trend expected of clock/bmal suppressors. (**F**) Putative *Sirt* expression shares this repressive trend. (**G-I**) Expression levels (DEseq2 norm) across time for all annotated genes identified by JTK cycle analysis reveal different numbers, amplitude, and timing of circadian gene cycles between young (36-119 days old) **(G)**, mid (2142-2146 days old) **(H)** and old (5869-5871 days old) **(I)** age groups. Plotlines represent an average of gene expression level as measured by LogCPM, each dot represents a specific sample, green-young (36-119 days old), purple-mid/adult (2142-2146 days old), and orange-old (5869-5871 days old). Tables S1A-S3 includes the lists of genes associated with these plots and heatmaps.

A comprehensive alignment of raw RNAseq *B. schlosseri* reads to *period* and *cryptochrome* alleles from diverse species (see methods for details), and a search for domains associated with both genes in the *B. schlosseri* genome also did not reveal putative homologous for these genes, suggesting that other *clock/bmal* regulators exist in tunicates.

After identifying key clock genes in the clock gene network, **we characterized their cyclical expression and found it varied widely with age.** We also identified genes with light-induced (entrained) expression (Table S1D), an important characteristic in circadian gene regulation. This was achieved through free-run and light pulse experiments which generated comprehensive, whole-organism transcriptional profiles from a total of 88 samples (multiple biological replicates, see annotations and gene counts in Table S1B-C and Methods). We identified clear, circadian expression of core-clock and clock-controlled genes using bioinformatics methods developed to find circadian *(18)* and chronologically differentially expressed genes (*16* and Methods) (Fig. 2, S2; Tables S2-3). The amplitudes and frequencies of these oscillations differed consistently between ages (Fig. 2G-I, S2; Tables S2-3). The number of genes that cycle throughout the day is higher in the mid-age group compared to old and young age groups (Mid>Old>Young). Each age group has unique sets of genes that cycle (Fig. 2C; Table S3). Both elderly and young colonies show a reduced amplitude of change and higher expression levels of clock-controlled genes. In middle-aged colonies, the amplitudes and frequencies of the gene oscillations are greater than their young and old counterparts (Fig. 2C).

**We studied cyclic gene dynamics at the tissue level by looking at the number of genes expressed at different time points in detailed tissue-specific gene profiles/enrichments***(15–16;* Table S4). This analysis revealed that the observed morphological and cellular aging phenotypes (Fig. 1) were accompanied by dramatic changes in the cycles of organ-specific gene expression (Fig. 3, Table S4).

**Fig. 3.**
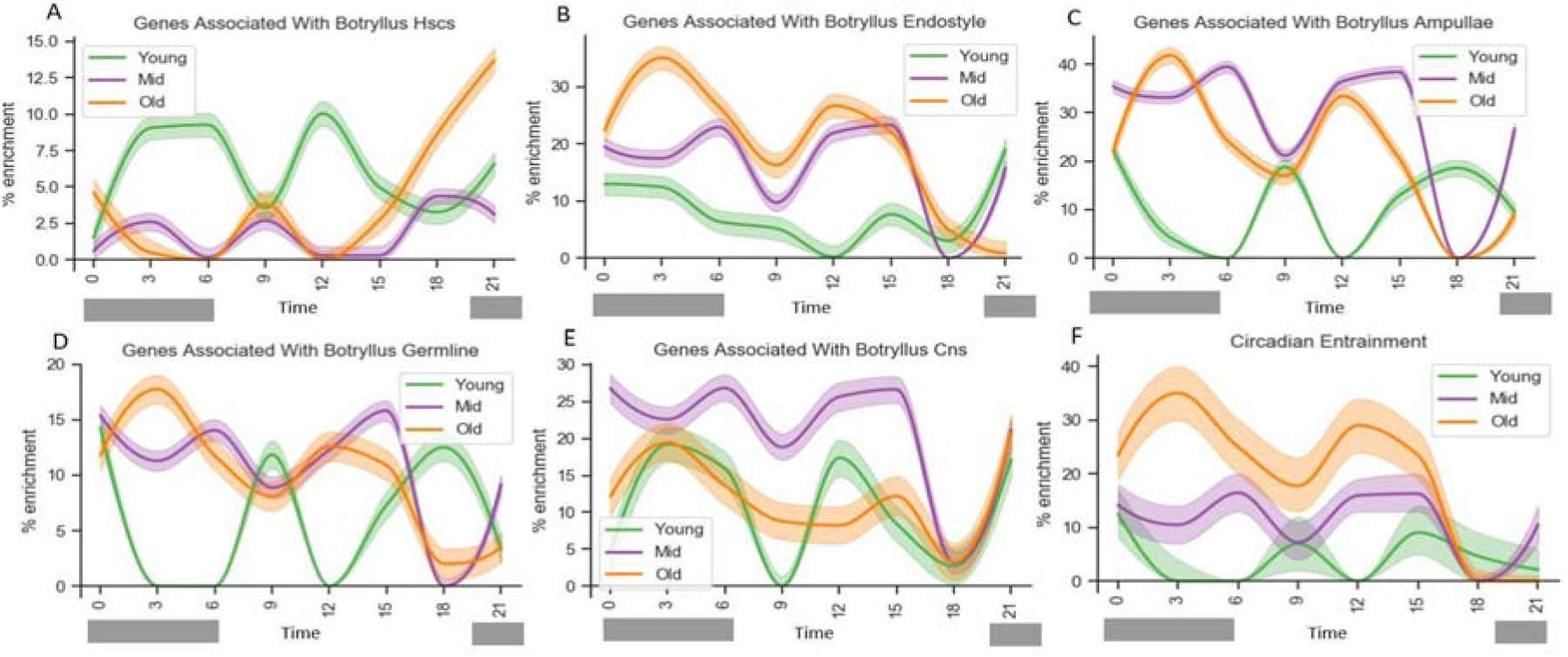
The percentage of genes dynamically expressed in tissues changes with age and correlates with the cycling dynamic of genes from the Circadian Clock Entrainment pathway. (A-E) Enrichment plots of tissue-specific gene signatures dynamic measured for 3 different age groups, every 3 hours over 24 hours (see methods). Percent of genes associated with (A) hematopoietic stem cells, (B) the endostyle stem cell niche, (C) blood vessels, (D) putative germline cells, (E) central nervous system, (F) circadian clock entrainment pathway expressed at each age and time point. (F) Enrichment plots of genes associated with the circadian entrainment pathway show age-specific oscillation patterns that correlate with the age and tissue-specific oscillation patterns observed in the majority of tissues and organs studied. Table S4 includes lists of the genes associated with these plots.

Significant age-associated changes were found in the cycling dynamics of genes associated with the hematopoietic system, the endostyle, the germline stem cells, and the central nervous system (Fig. 3 and Table S4). This is exemplified by the clear age-associated, cyclical enrichment pattern in the hematopoietic stem cell (HSC) gene profile. A higher percentage of HSC associated genes are dynamically expressed with larger, opposite amplitudes in young colonies compared to mid and old age groups (Fig. 3A). The endostyle associated genes behave differently, where throughout the day, the highest percentages of enrichment expression are observed in old colonies (Fig. 3B). This raised expression could be due to the old colonies attempting to compensate for reduced enriched HSC’s numbers in their endostyle niches *(13, 15)* (Fig. 1E). Young colonies also showed a diminished cycling pattern in the endostyle gene profile. The dynamic cycling of the gene enrichment associated with the *B. schlosseri* ampullae/blood vessels was similar to that of the endostyle. More genes were consistently expressed in old and mid-age colonies compared to the young. However, young colonies showed an undiminished cycling pattern that was the opposite of the pattern seen in mid and old colonies (Fig. 3C). The percent of genes associated with putative germline stem cell populations was higher in old and mid-age colonies. This resembled the ampullae enrichment patterns, where young colonies showed the near exact opposite cyclical patterns to mid and old colonies (Fig. 3D). The central nervous system (CNS) associated genes behaved differently than previous tissues discussed. In this system, the mid-age group showed the highest enrichment throughout the day. Unique to the CNS gene profile, all three age groups’ cycling dynamics were relatively similar (Fig. 3E). This is particularly intriguing considering the role of the CNS as the center of clock activity in mammals (specifically the suprachiasmatic nuclei)*(19)*. Investigating the specific genes associated with these patterns can point to CNS aging mechanisms by looking for deviations in the cycling dynamics of the mid-age group (Table S4).

This analysis revealed that tissue-specific gene expression oscillates in a daily cycle with different amplitudes and frequencies between ages. The number of genes that are expressed cyclically is generally higher in old and mid-age groups compared to the young. The age-associated tissue-specific oscillation patterns observed correlate with the age-specific oscillation patterns detected in genes associated with the circadian entrainment pathway (Fig. 3F). This demonstrates a strong link between clock-controlled genes and tissue activities.

After finding a link between aging and cyclic expression at the tissue level, we **studied the relationship between the circadian clock and aging in hundreds of known pathways**. This was made possible by having a comprehensive set of genes for each age and timepoint (Table S2A-C). These sets of genes were analyzed to find any associations to known pathways. We found over 450 pathways, all with some level of age-associated changes in their cycling dynamics (Table S5). Substantial circadian variation between age groups was discovered in the expression patterns of pathways linked to hallmarks of aging *(20–21)*, cellular processes, and tissue functions. (Table S5; Fig. 4). The age-specific patterns of these pathways (Fig. 4) correlated with the age-specific patterns of the circadian entrainment pathway (Fig. 3F). These cyclic patterns also correlated with the age-specific trends observed in the longevity regulation path (Fig.4P). These correlations indicate a strong link between the circadian clock and aging, suggesting that clock genes are key regulators of aging.

**Fig. 4.**
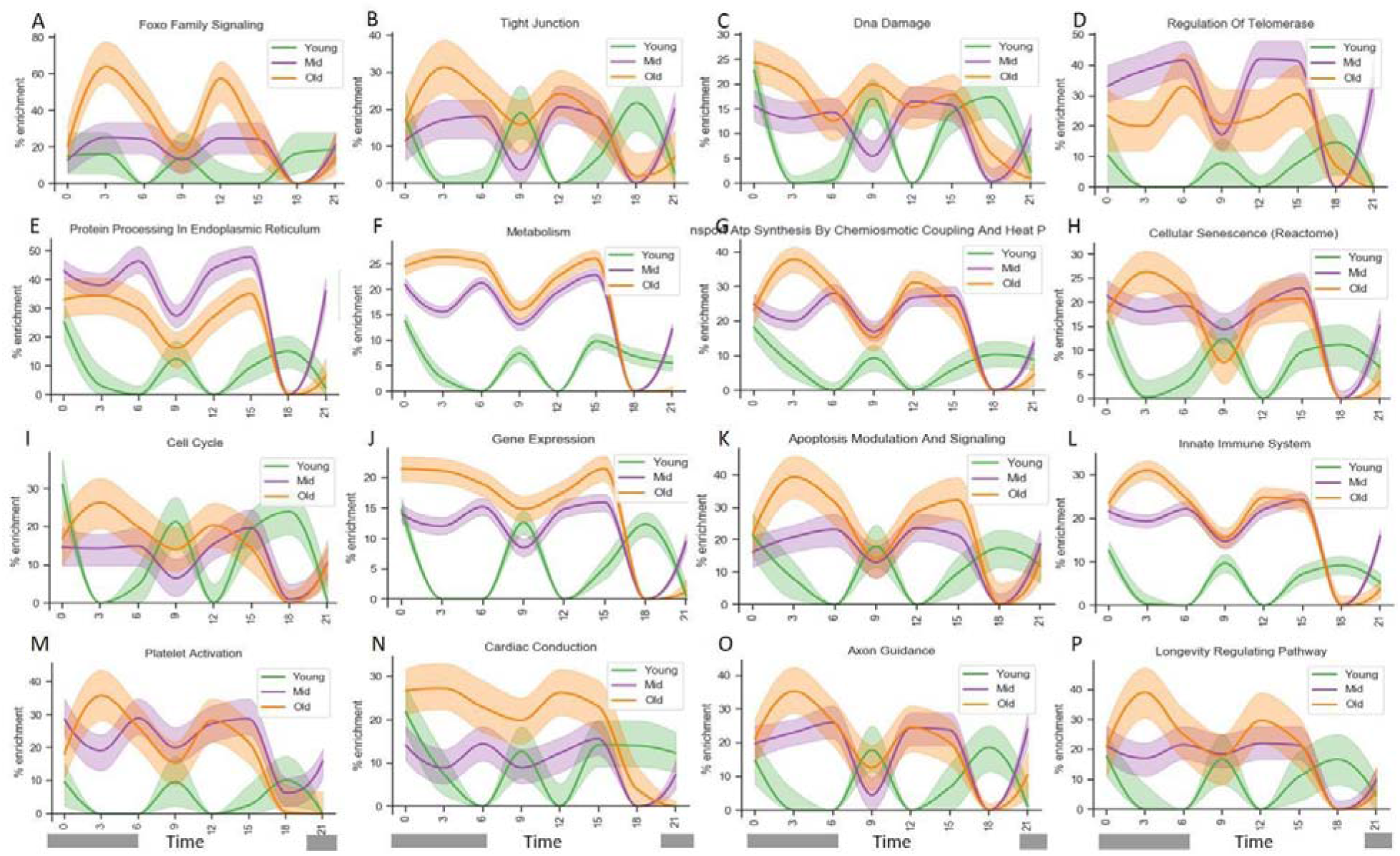
The cycling pattern of gene pathway profiles linked to hallmarks of aging, regulations of cellular processes, and tissue activities varied widely between ages. The effect of time and age on pathways associated with hallmarks of aging (*20*; A-H, P), cellular processes (I-K), and tissue activities (L-O) as measured by % of genes (from a specific pathway) activated at every time point across ages. (**A**) Stem cell exhaustion, Foxo signaling path *(21)*. (**B**) Altered intercellular communications, tight junction path. (**C**) Genomic stability, DNA damage path. (**D**) Telomere attrition, regulation of telomerase path. (**E**) Loss of proteostasis, protein processing in endoplasmic reticulum path. (**F**) Deregulated nutrient sensing, metabolism path. (**G**) Mitochondrial dysfunction, respiratory electron transport ATP synthesis path. (**H**) Cellular senescence path. (**I**) Cell cycle path. (**J**) Gene expression path. (**K**) Apoptosis modulation and signaling path. (**L**) Innate immune system path. (**M**) platelet activation path. (**N**) cardiac conduction path. (**O**) Axon guidance path. (**P**) Longevity regulating path.

Through building an aging-clock molecular atlas, this study generated the most complete transcriptomic database that links systemic changes in the molecular clock to aging to date. Using a model organism that is closely related to vertebrates with an intrinsic connection to stem cell aging *(22)*, this atlas emphasizes the extensive cycling variation in gene expression across ages, showcasing the importance that clocks must play in regulating stem cell aging.

## Acknowledgments

We thank C. Lowe, L. Crowder. C. Patton, J. Thompson, J. Lee, B. Compton, T. Naik L. Quinn for technical advice and help.

## Author contributions

Conception and design: A.V., Y.V., A.G., M.K., R.B., D.S.; mariculture, observation and sample collection: Y.V., A.G., K.J.I., K.J.P., A.V., C.A., B.R., T.G.,; RNA isolation and library preparation: K.J.P., A.V.; sequencing: N.F.N.; sequencing analysis and development of analytical tools: M.K., Y.V., A.G., D.S.,; flow cytometry: B.R.; writing of manuscript: A.V., A.G.,Y.V., M.K., R.B., I.L.W.; technical support and conceptual advice: A.V., R.B., D.S., N.F.N., S.R.Q., I.L.W.;

## Data and materials availability

RNA-Seq data are available on the Sequence Read Archive (SRA) database: BioProject: PRJNA682759

## Supplementary Figures

**Fig. S1:**
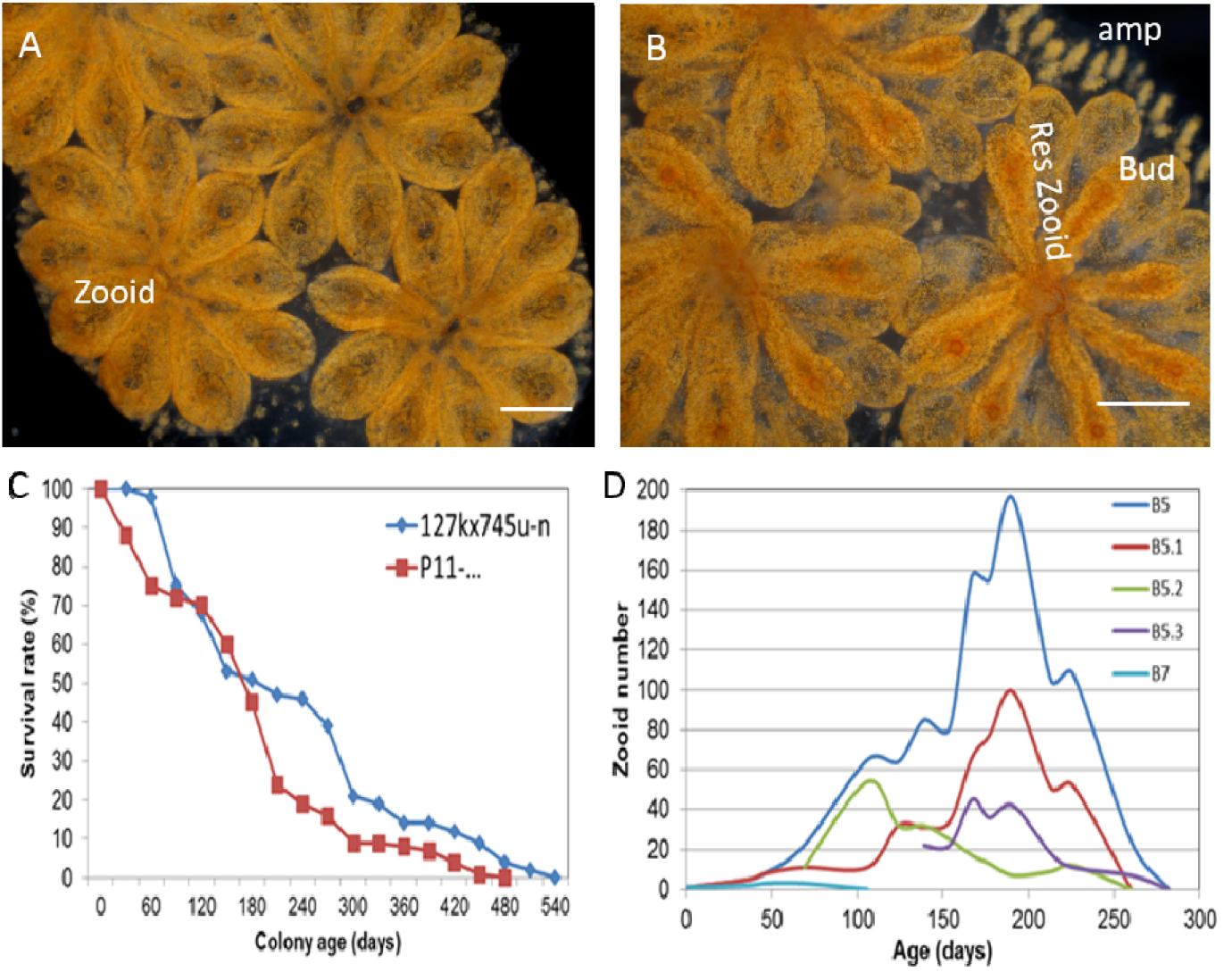
B. schlosseri asexual development, life span, and growth. *B. schlosseri* reproduces both through sexual and asexual (budding) pathways, giving rise to colonies (*A-B*). Upon settlement, the tadpole phase of the *B. schlosseri* lifecycle will metamorphose into a founder individual, which through asexual budding, generates a colony. *(**A-B**)* The colony includes three overlapping generations: an adult zooid, a primary bud, and a secondary bud, all of which are connected via a vascular network embedded within a gelatinous matrix (termed tunic) and terminate in finger-like protrusions (ampullae). This budding process continues weekly throughout the life of the colony producing multiple individuals (buds that grow into zooids) arranged in flower-shaped clusters called systems. (***A***) Bud development commences in stage A. Through budding, *B. schlosseri* generates its entire body, including digestive and respiratory systems, a simple tube-like heart, an endostyle that harbors a stem cell niche, a primitive neural complex, and siphons used for feeding, waste, and releasing larvae. Each week, successive buds grow large (***B***) and complete replication of all zooids in the colony, ultimately replacing the previous generation’s zooids, which die through a massive apoptosis. res zooid-resorbing zooid, *Scale bar-1mm (**C***) Survival plots of 162 colonies from two representative strains raised in our mariculture facility: 57 colonies in blue originate from the cross between colonies 127k and 745u-n and 105 originate from a cross between P11-1R and Bw221 (red). *B. schlosseri* colonies produce many offspring, ~25% die before reaching sexual maturity (2-3 months), about 50% die after 6 months and only ~10-20% survived over a year. (***D***) *B. schlosseri* colonies manifest stem-cell mediated tissue regeneration throughout adult life, which wanes with age (measured by number of individuals (zooids)) in a colony. The colony grows exponentially as a juvenile, reaching a maximum size near the last third of its life. Toward the end of its life, asexual reproduction begins to slow and the number of individuals in the colony is reduced because of budding failure (B5.1 B5.2 B5.3 are subclones of genotype B5).

**Fig. S2.**
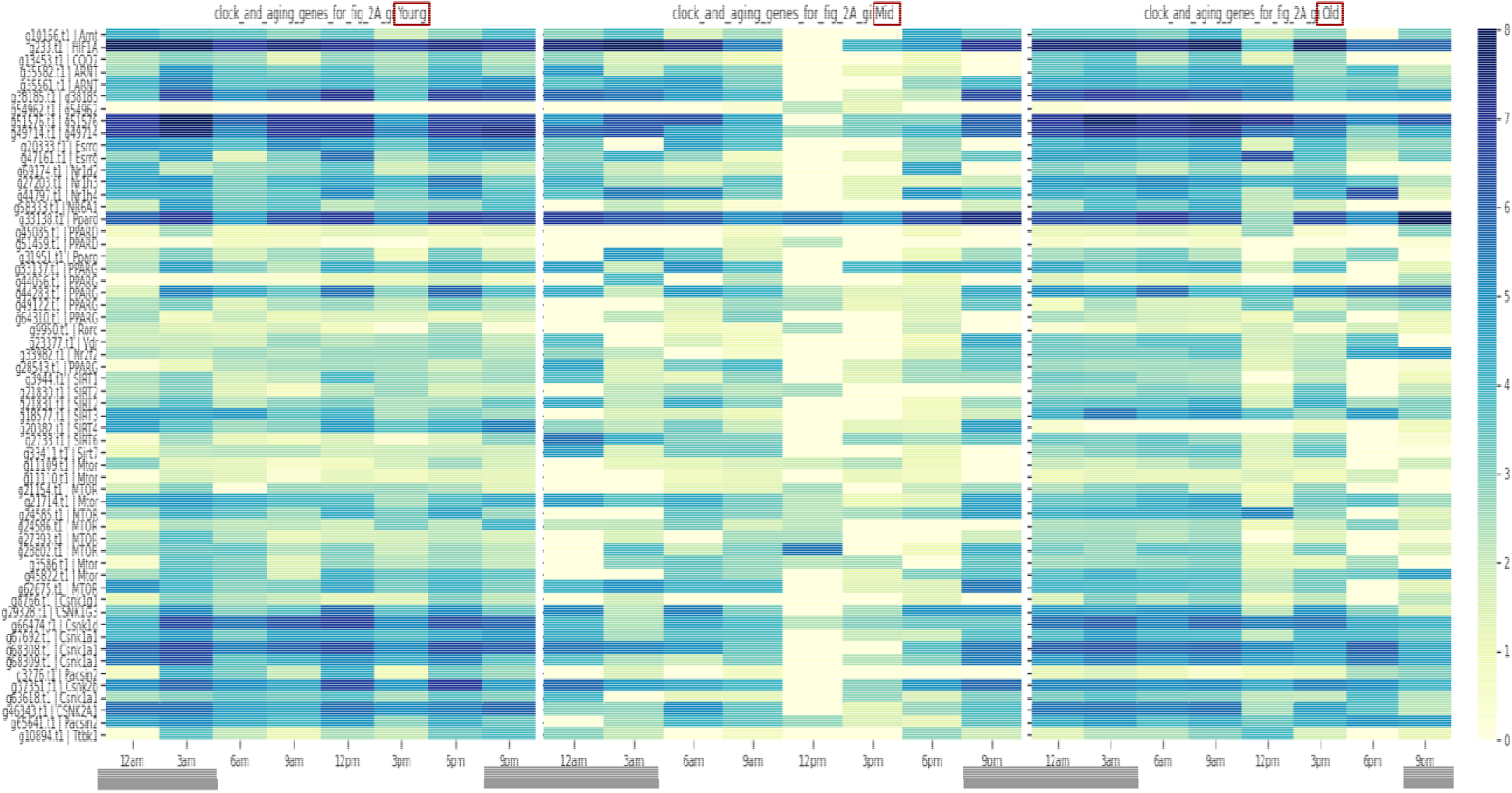
Age associated changes in *Botryllus* clock core gene cycling patterns. An average of gene expression levels (LogCPM) across time for putative clock and clock regulators genes. Different expression levels and timing of circadian gene cycles between young (36-119 days old; left), mid (2142-2146 days old; center) and old (5869-5871 days old; right).

## Materials and Methods

### Mariculture

Mariculture procedures have been described previously (*23*). Briefly, wild type *Botryllus schlosseri* colonies collected in Monterey bay, were tied to 3×5-cm glass slides and placed 5 cm opposite another glass slide in a slide rack. The slide rack was placed into an aquarium, and within a few days the tadpoles hatched, swam to the settlement slide, and metamorphosed into the adult body plan (oozooid). Single oozooids are then transferred to individual slides and grown at 18-20°C and under 14h/10h light/dark regimen (6am-8pm light/ 8pm-6am dark). Colonies were fed daily using a marine invertebrate diet prepared in the lab as described in Kowarsky et al. 2020).

### Identification of *B. schlosseri* aging phenotypes

Pictures of young and old colonies were taken. Morphological changes were recorded (e.g. pigmentation level, blood vessels shape). Zooid sizes were measured using image J (mm^2^). T-test was used to find significant changes.

### Recording diurnal behavior

We monitored heartbeat (per 30 sec) and siphon openings in zooids on developmental stage A1 during day (6pm light) and night (12am dark). Three ages were observed: 40 days old, (light n=12, dark n=10), 1515 days old, (light n= 9, dark n=6), and 6905 days old (light n=9 dark n=9). T-tests were used to find significant changes.

### Flow Cytometry

All flow cytometry analyses were performed using BD ACCURI-C6. Colonies were taken at two different ages: juvenile (3-5 days old; N=190) and old (4-13 years old, N=37). Cell suspension was isolated as described in *15*; briefly, *B. schlosseri* systems were meshed and filtered through a 40 μm mesh using a sterile 1 ml syringe pump. Cells were washed and collected in staining media: 3.3x PBS, 2% FCS and 10 mM Hepes. Cells were stained with 1) Propidium Iodide (PI), to differentiate live vs dead cells 2) alkaline phosphatase substrate measuring alkaline phosphatase activity in green fluorescence, 3) Cd49d-PE-Cy7, 4) CD57-Pacific Blue, 5) ConcavilinA-AF633, and 6) anti BHF (*25*) labeled with mouse serum APC-Cy7. Live cells were gated on a forward scatter (FSC) and side scatter (SSC) panels using log scale, and then gated further into different populations. Analysis of flow cytometry data was accomplished using FlowJo V10 (FlowJo).

### Clock experiments

#### Free-run experiment

Colonies from 3 different age groups (36-140 days old days; 2142-2146 days; 5869-5871 days) were placed in 1.5 liter tubs, 2-3 slides per tub and kept in dark. Following a minimum of 3 days under dark, whole systems (Fig. S1A) were sampled every 3 hours along the clock under red light: 12am (n=12), 3am (n=9) 6am (n=9), 9am (n=10), 12pm (n=10), 3pm (n=14), 6pm (n=11), 9pm (n=8). All samples were taken at either developmental stage A1 or A2 (Figure S1A).

#### Light pulse

Following 4 days under dark, samples (whole systems) were collected at midnight. 3 samples were exposed to light for half an hour (light pulse) and sampled afterwards. 5 colonies (100-135 days old; developmental stage A1-A2) were sampled in this experiment.

### Library preparation

All samples collected for RNAseq library prep (Detailed annotation in Table S1) were frozen in liquid nitrogen, and held at −80. RNA was prepared from frozen samples using Zymo Research Quick RNA MIcro Prep Kit #R1050, and cleaned using Zymo Research RNA Clean and Concentrator, #R1015. Samples were analyzed on an Agilent QC 2100 Bioanalyzer to determine quality prior to library preparation. cDNA was prepared using the Nugen Ovation RNA Sequencing System V2, #7102 and cleaned using the Qiagen QIAquick PCR purification kit, #28104, and then analyzed on the Agilent QC Bioanalyzer. If needed, samples were size-selected using Zymo Research Select-A-Size DNA Clean and Concentrator #D4080 prior to barcoding. Final library was prepared using NEB NEBNext Ultra II DNA Library Prep Kit #27645 and barcoded using NEBNext Multiplex Oligos for Illumina #E6609S. All magnetic bead purification was accomplished using BullDogBio CleanNGS RNA and DNA Spri Beads #CNGS005. Samples were then analyzed on the Agilent QC 2100 Bioanalyzer to determine the concentration of each sample prior to determining dilution prior to sequencing. On average, 12 million 2×150 bp reads (Illumina Nextseq 500) were sequenced for each library.

### Gene counts

Following sequencing, reads were processed using a Snakemake (***24***) pipeline: they were trimmed to remove low quality bases and primers, merged if the reads from both ends overlapped, and aligned to a database of *B. schlosseri* transcripts using bwa (mem algorithm), with likely PCR duplicates removed and then read counts determined for each transcript, resulting in a count matrix (Table S2).

### Gene orthology and methods used to search for cry and per

Gene orthology was based on sequence similarities between the *B. schlosseri* gene models and human and mouse gene annotations (BLAST score smaller than 10^−10^ as described in the *B. schlosseri* genome papers *11,25*). All discussions regarding putative clock genes are based on sequence and domains similarities alone. To search for the missing *period* and *cryptochrome* genes all alleles for both genes were taken from fly (flybase.org), fish (zfin.org), and frog (xenbase.org) databases and blasted against the *Botryllus* assembly using NCBI BLAST. Then all the alleles were combined to make a mini-assembly of these missing genes and aligned to the *B. schlosseri* RNA-seq data using Krypto and BWA-mem and the alignment was visualized with Jbrowse. A consensus sequence of the aligned RNA was looked for using Samtools, but none was found. Finally, we identified all the conserved domains in the Botryllus genome using NCBI domain search and focused on the ones that contained conserved domains that appear in *Per* and *Cry*. We generated a Maximum Likelihood Bootstrap Phylogenetic Tree with alleles and genes we thought were possible candidates using Clustal Omega Multiple Sequence Alignment and MegaX but did not identify putative *cry* or *per*.

### Identification of genes entrained by light

Differentially expressed genes between control colonies (kept in dark) and colonies that were exposed to light pulse at midnight were found using edgeR ((FDR < 0.05); *26* Table S1D).

### Identification of cycling genes

To identify circadian cycling genes we used MetaCycle (*18*), a package combining 3 different methods, (JTK_CYCLE, ARSER, Lomb-Scargle) developed to identify cycling genes. Each age group was tested by itself and all cycling genes identified by these programs are presented in Table S3A-C.

### Identification of time dependent gene signature and formation of binary tables

Identification of chronologically differential expressed genes and formation of binary tables have been described previously (*16*). Briefly, we used DEseq2 (*27*; FDR < 0.05) to identify differentially expressed genes between all possible combinations of contiguous and individual time points, resulting in a hierarchy of expressed time points for each gene for each age group. Based on these analyses, a binary gene-time expression matrix for every expressed gene recorded along the time points was produced for each age group individually, with the binary pattern chosen being the one with the highest correlation to the time hierarchy for that gene. With 1 indicating dynamically “high” expression and 0 indicating “low” or zero expression (Table S2A-C).

Using the binary matrix we identified pathways and Go terms associated with each age group for every time point (GeneAnalytics; *28;* Fig 4; Table S5).

### Gene enrichment plots

Gene enrichment plots have been described in *16*. Briefly, at each time point the proportion of genes in a gene set that are active (indicated by a 1 in the gene-time expression binary matrix defined above) is calculated. This gives a value between 0% (no genes in common) and 100% (all genes in the gene set are active at that time). A baseline expectation of the proportion of overlapping genes is calculated using a hypergeometric model that gives the likelihood that the same number of genes as in the selected gene set would be randomly selected from the matrix. In addition, the 68% confidence interval (1 standard deviation) of proportion of shared genes (‘enrichment’) from the hypergeometric is calculated and plotted, presented as a shaded region in the plot. Then the baseline is subtracted from the values calculated, with the confidence interval also subtracted, to show the expected range of values and how far the actual enrichment result differs from a null result. If the baseline expectation is greater than the actual enrichment (where the subtracted value would be negative) a value of 0% is used (as a negative percent was considered meaningless).

### Tissue enriched and specific pathways gene set used

For the gene enrichment plots the following gene sets were used:

1. For tissue specific nervous system, endostyle, ampullae, enriched hematopoietic stem cells (HSCs) and putative germ cells, we used existing *Botryllus* tissue enriched gene lists (*15, 16*; Table S4).
2. For the pathway and go terms identified by GeneAnalytics within our time data, we focused on the gene sets of the same names from PathCards. From each of these gene sets we removed any gene that did not have a putative homolog to a known *Botryllus* gene. As such the percentages within the enrichment plots refer to this curated *Botryllus* specific gene sets. If a gene name appeared more than once in the *Botryllus* gene model annotation (*11, 25*) all matching *Botryllus* gene ids were included in the gene set.

## Supplementary Tables

S1A *B. schlosseri* candidate clock core and regulators genes.

S1B Annotation of the 88 samples sequenced.

S1C Transcript counts for all 88 samples sequenced.

S1D Differentially expressed genes between control colonies (kept in dark) and colonies that were exposed to light pulse at midnight.

S2A-C A binary gene-time expression matrix for the 3 age groups.

S3A-C Meta cycle results for the 3 age groups.

S4 Tissue specific gene enrichment at every time point across the age groups (nervous (A-C), HSCs (D-G), endostyle (H-J), GSC (Q-T)).

S5 Pathway enrichment expression at every time point across ages based on GeneAnalytics (*28*) analysis superpaths enriched in Young (A), Mid (B) and Old (C) and the gene associated with them per every time point. (D) Color coded comparison between pathways enriched in all three ages.

Movie S1-Siphons opening during dark (night) versus light (day) of a young colony.

Siphon activity recorded at night versus day in young colonies, suggesting that *B. schlosseri* is a diurnal organism.

